# The Lomb-Scargle periodogram-based differentially expressed gene detection along pseudotime

**DOI:** 10.1101/2024.08.20.608497

**Authors:** Hitoshi Iuchi, Michiaki Hamada

## Abstract

**Motivation:** In recent years, single-cell RNA sequencing (scRNA-seq) has provided high-resolution snapshots of biological processes and has contributed to the understanding of cell dynamics. Trajectory inference has the potential to provide a quantitative representation of cell dynamics, and several trajectory inference algorithms have been developed. However, the downstream analysis of trajectory inference, such as the analysis of differentially expressed genes (DEG), remains challenging.

**Results:** In this study, we introduce a Lomb-Scargle (LS) periodogram-based algorithm for identifying DEGs associated with pseudotime in a trajectory analysis. The algorithm is capable of analyzing any inferred trajectory, including tree structures with multiple branching points, leading to diverse cell types. We validated this approach using simulated data and real datasets, and our results showed that our approach was superior when performing DEG analysis on complex structured trajectories. Our approach will contribute to gene characterization in trajectory analysis and help gain deeper biological insights.

**Availability:** All code used in our proposed method can be found at https://github.com/hiuchi/LS.

**Supplementary information:** Supplementary data are available at *Journal Name* online.

## Introduction

Single-cell RNA sequencing (scRNA-seq) provides high-resolution snapshots of biological processes and allows the observation of cellular state transitions, such as cell differentiation, cell cycle, and stimulus-response, at the single-cell level (Luecken and Theis (2019)). Indeed, trajectory inference, which is a computational technique based on single-cell transcriptomes for predicting biological processes, has advanced our understanding of cell dynamics across multiple disciplines, such as neuronal development (Bilgic *et al*. (2023) and Trevino *et al*. (2021)), immune cell differentiation (Le *et al*. (2020)), and state transitions in pathogenic organisms (Briggs *et al*. (2021)). These advances have been supported by bioinformatics.

To date, bioinformatic efforts have focused on developing trajectory inference algorithms (Saelens *et al*. (2019)). Most trajectory inference approaches involve the following steps: (1) to reduce the dimensionality of the gene × cell matrix; (2) to assign each cell to a lineage in the reduced space; and (3) to assign a pseudotime based on the distance of each cell from the starting point. Many trajectory inference algorithms have been developed with variations in dimensionality reduction and lineage assignment (Saelens *et al*. (2019)). Despite the active development of trajectory inference, methodologies for downstream analyses are lacking.

Essential downstream trajectory inference analysis is used to identify differentially expressed genes (DEGs). In time-series omics data analysis, there are two patterns of DEGs: one is a statistical test of whether a gene is dynamically expressed along pseudotime, and the other is a test of whether a gene’s expression is shifted in the two conditions (e.g., wildtype vs. knockout). In this paper, the former is referred to as the dynamic gene expression test, and the latter as the shifted gene expression test. Although the dynamic gene expression test is the mainstream test developed for time-series omics data analysis, shifted gene expression is also important for characterizing changes in gene expression patterns by comparing wildtype and knockout and by the presence or absence of stimuli. In this study, we aimed to develop an algorithm that can perform both dynamic and shifted gene expression tests.

Additionally, the development of scRNA-seq experiments and analysis methods generates complex trajectories, such as tree structures with branches, and there is a need for novel downstream analysis methods to address this dataset. Typical downstream analyses include TradeSeq (Van Den Berge *et al*. (2020)), PseudotimeDE (Song and Li (2021)), and Lamian (Hou *et al*. (2021)). All these are model-based approaches, representing gene expression as a function of pseudotime and computing statistics such as P-values. Although the model-based approach is powerful and widely used, its drawback lies in the difficulty of expressing branching data mathematically. Thus, the maturation of single-cell transcriptome experiments and trajectory inference require new approaches.

This study introduces a Lomb-Scargle (LS) periodogram-based algorithm for identifying DEGs associated with pseudotime. Although the fast Fourier transform, which is often used in time-series analysis, requires uniform sampling points, it is impossible to control the pseudotime precisely. Here, the Lomb-Scargle periodogram can transform time-series data with non-uniform sampling points into frequency-domain data. Our approach involves transforming pseudotime domain data from scRNA-seq and trajectory inference into frequency-domain data using LS. This versatile method is capable of analyzing any inferred trajectory, including tree structures with multiple branching points, leading to diverse cell types. We validated this approach using simulated and real datasets, demonstrating its efficacy as a robust, model-independent tool for trajectory analysis.

## Method

A schematic diagram of the proposed method is shown in Figure 1. In this scenario, the gene expression of the WT is branched, which is difficult to model as a function; however, the proposed method can analyze it, even if it is branched or a combination of branching and unbranching.

**Fig. 1.**
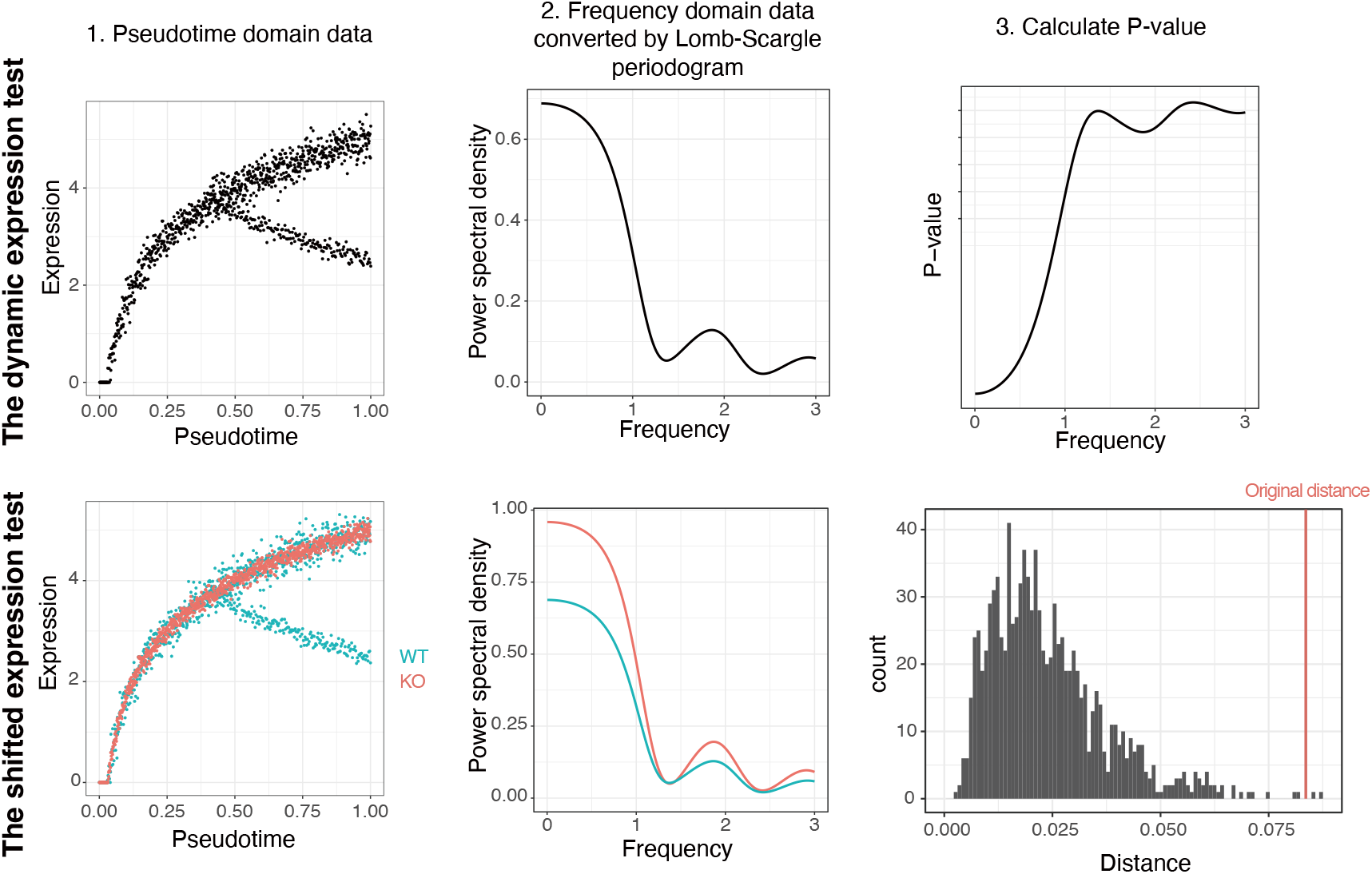
Procedure of the proposed method. The top row is the dynamic expression test flow, and the bottom row is the shifted expression test. Our approach can also fairly judge divergent gene expression, as in this example. In both tests, the pseudotime domain data are first converted into frequency domain data by Lomb-Scargle. In the dynamic expression test, the P-value is calculated for each frequency, and the smallest P-value is considered as the P-value of the gene. If the P-value of a frequency is below the significance level, the gene is considered to be significantly dynamic. In the shifted expression test, the pseudotime of the original expression data is shuffled to obtain the null distribution, and the P-value is calculated. In this case, 1000 permutation tests were performed, and a distance longer than the original distance was obtained twice, so 2/1000 = 0.002 is the P-value.

### Definition of the Lomb-Scargle periodogram

We provide an overview of the Lomb-Scargle periodogram (Leroy (2012) and VanderPlas (2018)). Given a set of *N* gene expressions *y*_*i*_ at the observed pseudotime *t*_*i*_ (*i* = 1, …, *N*), the Lomb-Scargle periodogram at frequency *f* is defined as:

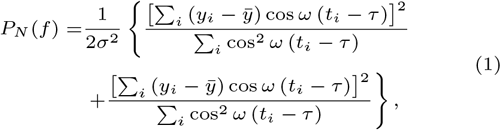

where 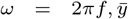 and *σ*^2^ are the mean and variance, respectively, given by

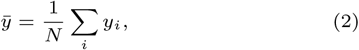

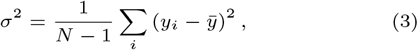

and the time offset *τ* is defined as:

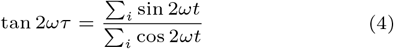

See VanderPlas (2018) for the derivation of the Lomb-Scargle periodograms and the details.

### The dynamic gene expression test

Pseudotime domain gene expression data are transformed into the Frequency domain by the formula (1). The smallest P-value among the P-values of each power spectrum in the frequency domain was considered the P-value of the gene. In other words, if a gene has a significant power spectrum, our method regards it as being associated with pseudotime.

### The shifted gene expression test

In this test, the expression of gene *g* under the two conditions is first converted into frequency-domain data by the formula (1). The distance (Euclidean or Canberra) *D*_*g*_ between the two frequency domains is calculated. The Canberra distance between the two vectors *p* and *q* is calculated as

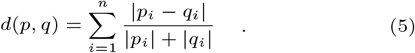

The P-value is calculated using the permutation test. First, to obtain the null distribution *N*_*g*_, the pseudotime of gene *g* was shuffled (e.g., 1000 times), and the distances were calculated after the domain transformation. *Dg > Ng* was counted and divided by the number of shuffles (1000 time here) to obtain the P-value.

In the shifted gene expression test, we calculated the distance between the frequency domain vectors from two conditions (wildtype and knockout) and determined a P-value based on the actual distance’s position within the null distribution.

### Simulation data synthesis and trajectory inference

In brief, simulation data for the performance tests were generated using the dynverse framework (Saelens *et al*. (2019)) to synthesize the count data, after which the trajectory and pseudotime were estimated using Monocle3 version 1.3.1 (Trapnell *et al*. (2014)) and Slingshot version 2.0.0 (Street *et al*. (2018)). All count data were generated using dyno version 0.1.2 or dyntoy version 0.9.9. The count data for Figure 2 were generated using the generate_dataset(model=“linear” or “bifurcating,” num cells=250 or 500 or 1000, num_features=1000, differentiall _expressed_rate=0.5) function in the dyntoy package. These data include the correct label as to whether it is a DEG and the pseudotime assigned to each cell.

**Fig. 2.**
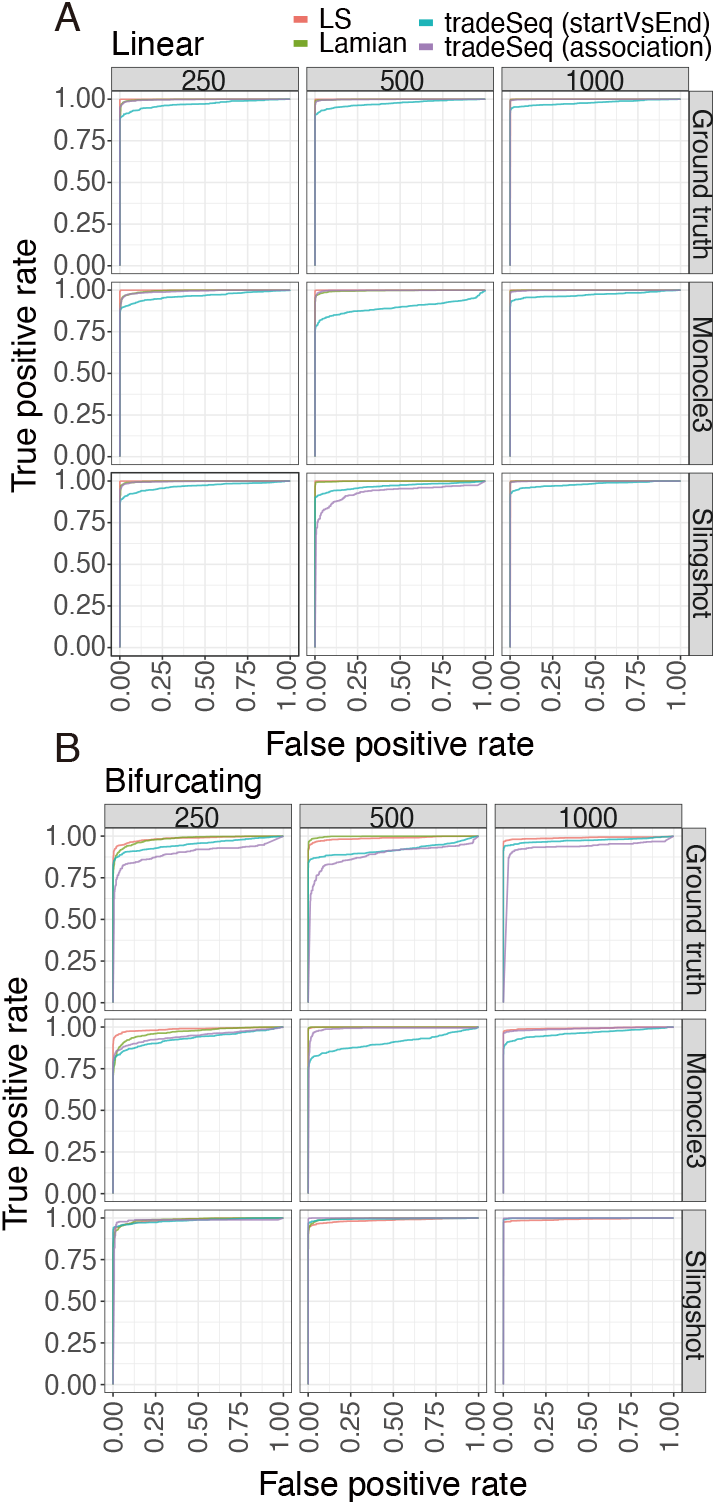
The benchmark of the dynamic expression test using synthetic data. Performance was compared using the ROC curve. (A) Linear and (B) bifurcating trajectories were generated by the dyneverse package (Saelens *et al*. (2019)). DEG analysis was performed on the ground truth trajectory, which is the correct trajectory, and on trajectories estimated by two types of trajectory inference. The colors indicate the DEG analysis algorithm. The numbers above the boxes indicate the number of cells. The origin of the pseudotime is shown to the right of the box. ROC, receiver operatorating characteristic; DEGs, differentially expressed genes.

The simulation data shown in Figure 3 were synthesized using the dyno package. Initialise_model, generate_tf _network, generate_featur _network, generate_kinetics, and generate _gold_standard function were used to simulate the gene expression of the wildtype and C1_TF1 gene knockout strains. Simulations were performed for 1000 cells and sparse data were generated by random sampling from the original data (see Supplementary Information for detailed procedures).

**Fig. 3.**
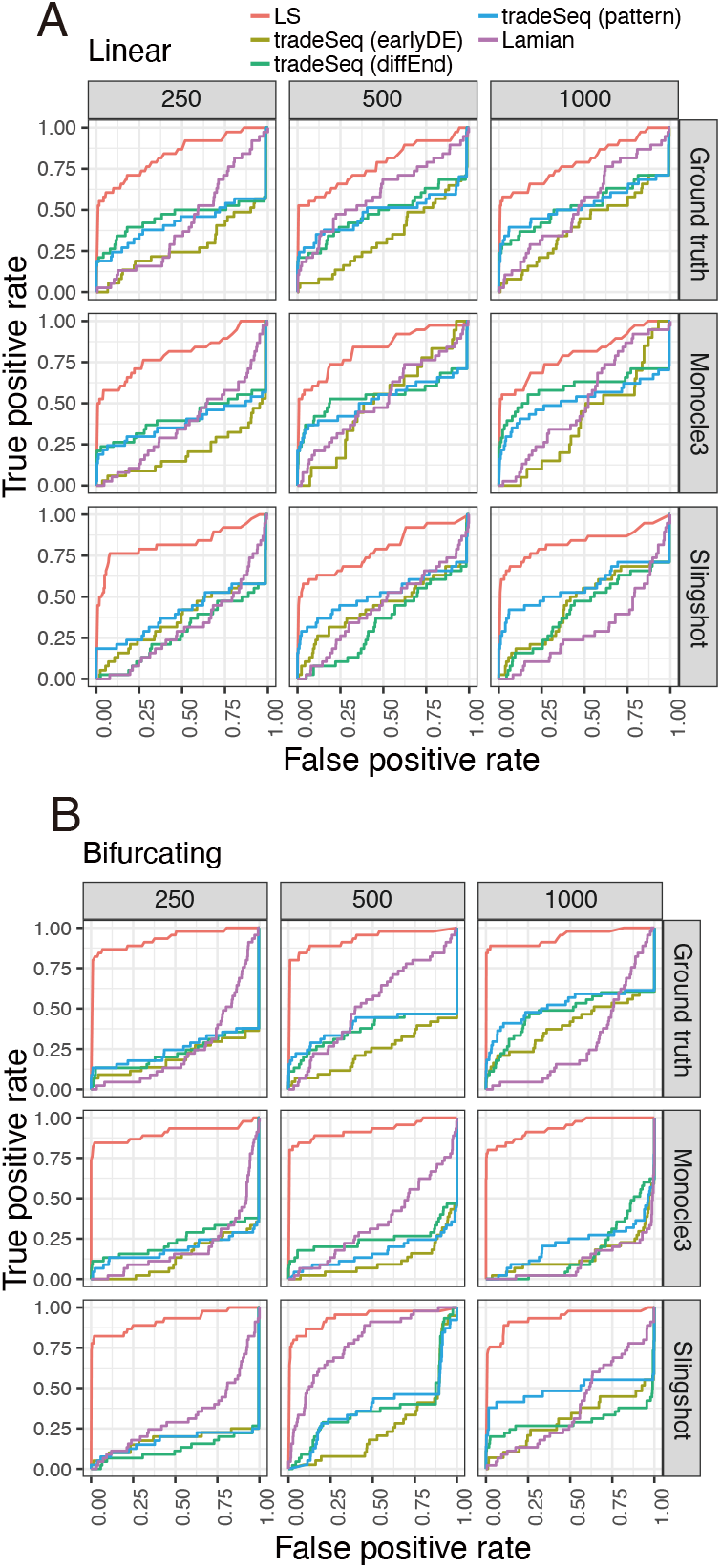
The benchmark of the shifted expression test using synthetic data. Performance was compared using the ROC curve. (A) Linear and (B) bifurcating trajectories were generated by the dyneverse package (Saelens *et al*. (2019)). Two simulated trajectories (WT and KO) were synthesized for each comparison to test whether the two trajectories were statistically differentially expressed. DEG analysis was performed on the ground truth trajectory, which is the correct trajectory, and on trajectories estimated by two types of trajectory inference. The colors indicate the DEG analysis algorithm. The numbers above the boxes indicate the number of cells. The origin of the pseudotime is shown to the right of the box. ROC, receiver operating characteristic; DEG, differentially expressed gene; WT, wildtype; KO, knockout.

**Fig. 4.**
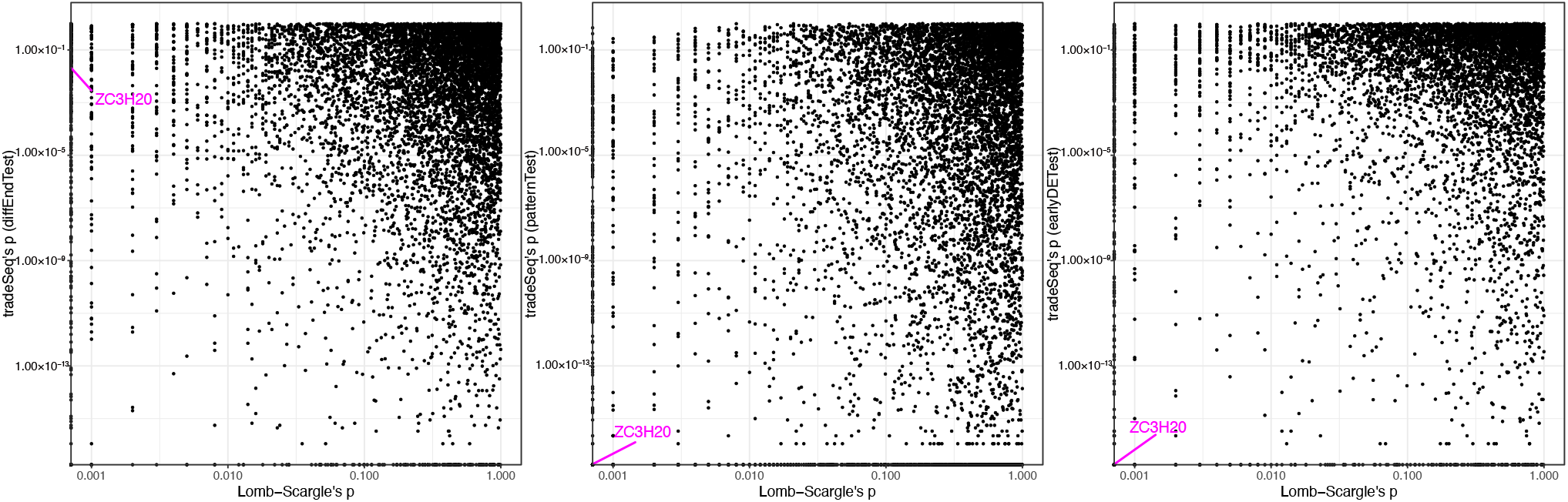
Comparison of P-values of the shifted test between the proposed method and three tests of tradeSeq (diffEndTest/patternTest/earlyDETest) on a Trypanosoma single-cell RNA sequencing (scRNA-seq) dataset. The authors of the original paper aimed to understand the transition mechanisms between replicative “slender” and transmissible “stumpy” bloodstream types of trypanosomes. They performed scRNA-seq analysis of the WT and ZC3H20 KO strains, an important state transition factor. Pre-processed data, including trajectory analysis by Slingshot, was kindly provided by Ross Laidlaw, University of Glasgow (Laidlaw *et al*. (2022)). As ZC3H20 is knocked out, a small P-value is expected.

Additionally, to test the robustness of each method, a DEG decision was made using pseudotime estimated using Monocle3 and Slingshot. The trajectory and pseudotime estimation by Monocle3 was performed by the following functions (specified arguments in parentheses), preprocess_cds(num_dim = 50), reduce dimension(preprocess_method = “PCA”, reduction_method = “UMAP”), cluster_cells(), learn_graph(), order cells (the argument root cells was the cell set as root during the simulation). For the estimation of trajectory and pseudotime using Slingshot, the expression levels of each cell were normalized to have the same distribution and then logarithmized. Clustering was performed using Gaussian mixture modeling with mclust version 6.0.0 using PC1 and PC2 obtained using the prcomp function. Using these cluster labels and PCA planes, the trajectory and pseudotime were calculated using the slingshot function.

### DEG detection by comparative methods

These analyses were performed using default parameters unless otherwise stated.

#### tradeSeq

The tradeSeq version 1.7 uses a negative binomial generalized additive model framework to model gene expression for each lineage. Modeling was performed using the fitGAM function (nknots=6). For comparisons between two groups, each group was analyzed as a lineage. The created models were evaluated using the associationTest, startVsEndTest, diffEndTest, patternTest, and earlyDETest (knots = c(1, 3)) functions to obtain P-values.

#### Lamian

Lamian version 0.99.1 is a framework that implements statistical analysis for various pseudotime analyses. We used the lamian_test(permuiter = 100) function with est.type=“chisq” and test.type=“time” for determining whether gene expression was associated with pseudotime and for comparing the two groups, respectively.

### Comparison of computation time

The computation times for each method were compared using an iMac (24-inch, M1, 2021) with a single core. Computation times were measured using the tictoc package, version 1.2. Simulation data were generated using the dyno and dyngen packages as previously described. The simulation data contained 2,000 genes, and datasets were generated for 100, 500, 1000, and 5000 cells.

### Practical analysis with a Trypanosoma dataset

Each method was evaluated using publicly available Trypanosoma scRNA-seq data (Briggs *et al*. (2021)). We aimed to understand the developmental process of Trypanosoma by knocking out the developmental regulator ZC3H20 and performing scRNA-Seq. The analyzed dataset, including two-dimensional projection maps and trajectory analysis using Slingshot, was kindly provided by Ross Laidlaw, University of Glasgow (Laidlaw *et al*. (2022)). DEG analysis of the dataset was performed using the proposed method and tradeSeq.

## Results

### Simulation dataset

Simulation data were used for benchmarking under controlled conditions (Figure 2 and 3). Although these synthetic data were cleaner than the actual experimental data, they were valuable for performance comparisons. First, the simulation data were generated to verify whether a gene was dynamically expressed over pseudotime (Figure 2). We conducted benchmarking using not only the correct pseudotime based on the ground truth but also the pseudotime estimated by Monocle3 and Slingshot. Because the accurate estimation of pseudotime remains a difficult issue, we compared the performance with imprecise pseudotime, which is closer to the actual analysis (Figure 6 and 7).

### The dynamic gene expression test

First, simulated data were generated to assess the performance of each method in a dynamic test to determine whether each gene was dynamically expressed in pseudotime. Note that the pseudotime estimated by Monocle3 and Slingshot is subject to variation, which depends on the trajectory inference (Figure 6). To clarify the effect of each trajectory inference and cell number, the accuracies of the linear trajectory (Figure 2A and Table 1) and bifurcated data (Figure 2B and Table 2) were evaluated using receiver operatorating characteristic curve (ROC) by combining these conditions. The low-noise and clear synthetic data resulted in an area under the curve (AUC) close to 1 for most conditions, reflecting the fact that Monocle3 was less accurate at predicting pseudotime than Slingshot. Lamian did not work properly under certain conditions and returned errors; it is described as an NA in Table 2. In summary, the methods that functioned successfully demonstrated high performance regardless of the cell number or trajectory inference.

**Table 1.**
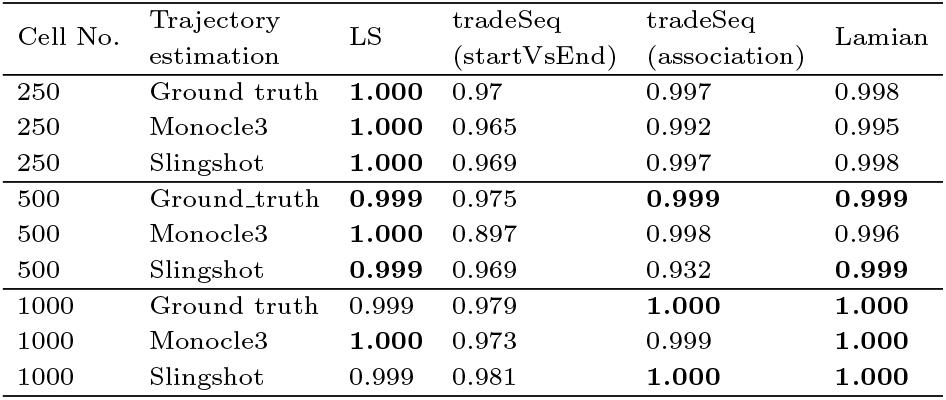
Area under the curve of Figure 2A.

**Table 2.** Area under the curve of Figure 2B.

| Cell No. | Trajectory estimation | LS | tradeSeq (startVsEnd) | tradeSeq (association) | Lamian |
| --- | --- | --- | --- | --- | --- |
| 250 | Ground truth | **0.985** | 0.951 | 0.899 | 0.982 |
| 250 | Monocle3 | **0.987** | 0.932 | 0.945 | 0.969 |
| 250 | slingshot | 0.988 | 0.984 | 0.982 | **0.989** |
| 500 | Ground truth | **0.988** | 0.92 | **0.886** | **0.998** |
| 500 | Monocle3 | **1.000** | 0.908 | 0.988 | **1.000** |
| 500 | slingshot | **0.986** | 0.995 | **1.000** | **0.997** |
| 1000 | Ground truth | **0.990** | 0.974 | 0.93 | NA |
| 1000 | Monocle3 | **0.993** | 0.963 | 0.99 | NA |
| 1000 | slingshot | 0.992 | **0.999** | **0.999** | NA |

### The shifted gene expression test

Next, we assessed the detection accuracy of shifted gene expression under two conditions (WT and KO). The boxes show the actual P-values calculated using each method. Note that in this simulation scenario, the gene C1 TF1 is knocked out; therefore, expression was lost in the KO strain, and a small P-value was obtained for LS only. In this comparison, the proposed method outperformed the existing methods under all conditions. The accuracy of pseudotime estimation using Monocle3 and Slingshot is shown in Figure 7. In this dataset, the proposed method showed a high AUC despite the low accuracy of pseudotime estimation in Monocle3 and Slingshot (Figure 3 A, B, and Tables 3 and 4). These comparison methods were not designed to bifurcate data. The fact that the proposed method outperforms other methods on data for which predicting pseudotime is difficult indicates the robustness of our method.

**Table 3.** Area under the curve of Figure 3A.

| Cell No. | Trajectory estimation | LS | tradeSeq (diffEnd) | tradeSeq (pattern) | tradeSeq (earlyDE) | Lamian |
| --- | --- | --- | --- | --- | --- | --- |
| 250 | Ground truth | **0.836** | 0.457 | 0.423 | 0.279 | 0.418 |
|  | Monocle3 | **0.800** | 0.415 | 0.384 | 0.206 | 0.373 |
|  | Slingshot | **0.826** | 0.290 | 0.405 | 0.371 | 0.314 |
| 500 | Ground truth | **0.777** | 0.485 | 0.482 | 0.334 | 0.602 |
|  | Monocle3 | **0.833** | 0.550 | 0.528 | 0.509 | 0.529 |
|  | Slingshot | **0.787** | 0.345 | 0.519 | 0.448 | 0.440 |
| 1000 | Ground truth | **0.797** | 0.530 | 0.536 | 0.404 | 0.537 |
|  | Monocle3 | **0.802** | 0.602 | 0.525 | 0.406 | 0.519 |
|  | Slingshot | **0.830** | 0.430 | 0.565 | 0.463 | 0.307 |

**Table 4.** Area under the curve of Figure 3B.

| Cell No. | Trajectory estimation | LS | tradeSeq (diffEnd) | tradeSeq (pattern) | tradeSeq (earlyDE) | Lamian |
| --- | --- | --- | --- | --- | --- | --- |
| 250 | Ground truth | **0.934** | 0.232 | 0.252 | 0.204 | 0.260 |
|  | Monocle3 | **0.907** | 0.241 | 0.189 | 0.135 | 0.198 |
|  | Slingshot | **0.919** | 0.127 | 0.176 | 0.183 | 0.323 |
| 500 | Ground truth | **0.932** | 0.368 | 0.392 | 0.236 | 0.526 |
|  | Monocle3 | **0.923** | 0.249 | 0.164 | 0.109 | 0.349 |
|  | Slingshot | **0.939** | 0.369 | 0.381 | 0.250 | 0.785 |
| 1000 | Ground truth | **0.945** | 0.474 | 0.518 | 0.397 | 0.288 |
|  | Monocle3 | **0.939** | 0.161 | 0.216 | 0.143 | 0.104 |
|  | Slingshot | **0.933** | 0.297 | 0.489 | 0.307 | 0.389 |

### Practical analysis with a Trypanosoma dataset

Next, to evaluate each method using real experimental data, we compared each method on a Trypanosoma scRNA-seq dataset. The authors knocked out ZC3H20, which is a key factor in the cell form transition between the replicative slender form and transmissible stumpy bloodstream form, and performed scRNA-seq Briggs *et al*. (2021). Therefore, ZC3H20 expression was absent in the KO strain. DEG analysis of these data using the proposed method and tradeSeq (diffEndTest/patternTest/earlyDETest) showed very low P-values for ZC3H20, except for diffEndTest. This indicates that the proposed method can identify the genes that must be detected. In addition, the variation in P-values between methods indicated that the results of DEG analysis in the trajectory analysis varied from method to method. In addition, gene ontology analysis was performed using the significant genes from the dynamic and shifted expression tests (Tables 5 and 6). The results indicated that the terms enriched in the dynamic expression test included several ATP-related pathways. Another report by the same authors identified ATP-related genes as DEGs only in the transmissible stumpy bloodstream form (Briggs *et al*. (2023)). Additionally, the knockdown of glycolytic genes was involved in cell cycle arrest (Marques *et al*. (2022)), and the role of these genes in form transition is expected to be clarified in future studies.

**Table 5.** Significant gene ontology terms of dynamic expression test.

| Term ID | Term name | P-value |
| --- | --- | --- |
| GO:1904949 | ATPase complex | 6.67.E-04 |
| GO:0016469 | proton-transporting two-sector ATPase complex | 6.67.E-04 |
| GO:0015078 | proton transmembrane transporter activity | 1.84.E-03 |
| GO:0015318 | inorganic molecular entity transmembrane transporter activity | 2.14.E-03 |
| GO:0009141 | nucleoside triphosphate metabolic process | 2.58.E-03 |
| GO:0009199 | ribonucleoside triphosphate metabolic process | 2.58.E-03 |
| GO:0005929 | cilium | 2.81.E-03 |
| GO:0120025 | plasma membrane bounded cell projection | 2.81.E-03 |
| GO:0042995 | cell projection | 5.56.E-03 |
| GO:0070925 | organelle assembly | 8.53.E-03 |
| GO:0046034 | ATP metabolic process | 8.53.E-03 |
| GO:0009144 | purine nucleoside triphosphate metabolic process | 8.53.E-03 |
| GO:0009205 | purine ribonucleoside triphosphate metabolic process | 8.53.E-03 |
| GO:0007018 | microtubule-based movement | 1.23.E-02 |
| GO:0009142 | nucleoside triphosphate biosynthetic process | 1.29.E-02 |
| GO:0009201 | ribonucleoside triphosphate biosynthetic process | 1.29.E-02 |
| GO:0045259 | proton-transporting ATP synthase complex | 1.29.E-02 |
| GO:0022890 | inorganic cation transmembrane transporter activity | 1.35.E-02 |
| GO:0009206 | purine ribonucleoside triphosphate biosynthetic process | 4.99.E-02 |
| GO:0015986 | proton motive force-driven ATP synthesis | 4.99.E-02 |
| GO:0006754 | ATP biosynthetic process | 4.99.E-02 |
| GO:0003341 | cilium movement | 4.99.E-02 |
| GO:0009145 | purine nucleoside triphosphate biosynthetic process | 4.99.E-02 |
| GO:0140662 | ATP-dependent protein folding chaperone | 4.99.E-02 |
| GO:0099568 | cytoplasmic region | 5.00.E-02 |
| GO:0097014 | ciliary plasm | 5.00.E-02 |
| GO:0032838 | plasma membrane bounded cell projection cytoplasm | 5.00.E-02 |
| GO:0030684 | preribosome | 5.00.E-02 |

**Table 6.** Significant gene ontology terms of shifted expression test.

| Term ID | Term name | P-value |
| --- | --- | --- |
| GO:0005930 | axoneme | 2.13E-02 |
| GO:0044183 | protein folding chaperone | 3.17E-03 |
| GO:0043168 | anion binding | 4.50E-02 |

### Evaluation of runtime

Finally, we compared the computational costs of each method when the number of cells varied from 100 to 5000 (Figure 5) to test whether gene expression is associated with pseudotime. LS required the least computational time, whereas testing whether the genes shifted expression under the two conditions had the highest computational cost. Most of the computational time was spent on the permutation test to calculate P-values. Although the computational cost of all methods increased with the cell number, it was sufficiently less than the time spent on cell culture, animal experiments, and sequencing.

**Fig. 5.**
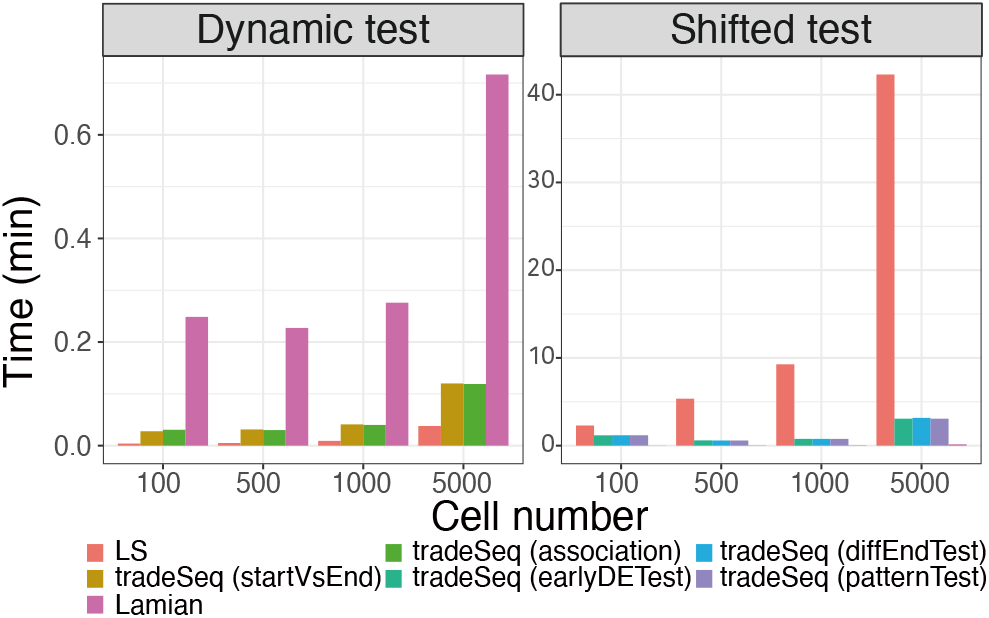
A comparison of the computation time of each method. The computational time to analyse the simulation data is measured and each dataset contains 2,000 genes.

## DISCUSSION

This study achieved highly accurate DEG detection in the scRNA-seq data by applying frequency-domain analysis (Figure 2 and 3). Time-series omics data from bulk RNA-seq and mass spectrometry are generally difficult to analyze in the frequency domain using fast Fourier transforms because the time points are limited, and the time periods are often short, as samples are collected at each time point by killing the animals. For circadian rhythm studies, where time-series data are frequently acquired, sampling every two hours for two days is recommended (Hughes *et al*. (2017)); however, experimental data following this guideline are rare owing to their high cost and burden. Thus, bulk omics time-series data have a small number of time points (a few to a dozen at most) and are sparse (sampled once every few hours) compared to classical time-series data such as electroencephalograms or electromyography. Therefore, the application of common time-series data analysis algorithms, such as fast Fourier transforms, has been limited, and their performance has been poor in practical cases (Iuchi *et al*. (2018)). Here, trajectory inference can be considered high-quality time-series data with thousands or tens of thousands of points by considering each cell as a single time point. This technological breakthrough has made it possible to apply

Fourier transforms and Lomb-Scargle periodgrams to biological time-series data. Thus far, there have been few examples of frequency-domain analysis applied to trajectory inference, and more applications are expected in the future.

The advantage of our proposed method is that it is a generic approach that is not limited by the trajectory structure. In complex biological phenomena, the trajectory structure may differ between WT and KO mice; for example, the trajectory may diverge into multiple branches or branch multiple times. It is difficult to model such complex structured trajectories and compare them in two groups; however, by transforming them into the frequency domain, any structured trajectory can be reduced to a vector-to-vector comparison problem. On the other hand, the model-based approach also has its advantages. If gene expression in the early phase of pseudotime is not important but is important in the late phase, the diffEndTest of tradeSeq is the most promising candidate. As previously stated, the critical issues in DEG analysis are defining the genes to be extracted and selecting the appropriate DEG analysis algorithm (Iuchi and Hamada (2021)). The results show that our approach is superior when performing DEG analysis on complex trajectories.

A unique feature of this study is the comparison with not only the ground truth pseudotime but also the pseudotime estimated by Monocle3 and Slingshot (Figure 2, 3, 6 and 7). These experiments revealed the influence of pseudotime on the results of DEG analysis. The combination of trajectory inference and DEG analysis algorithms is an important issue that needs to be optimized in the future through comprehensive and fair comparisons.

**Fig. 6.**
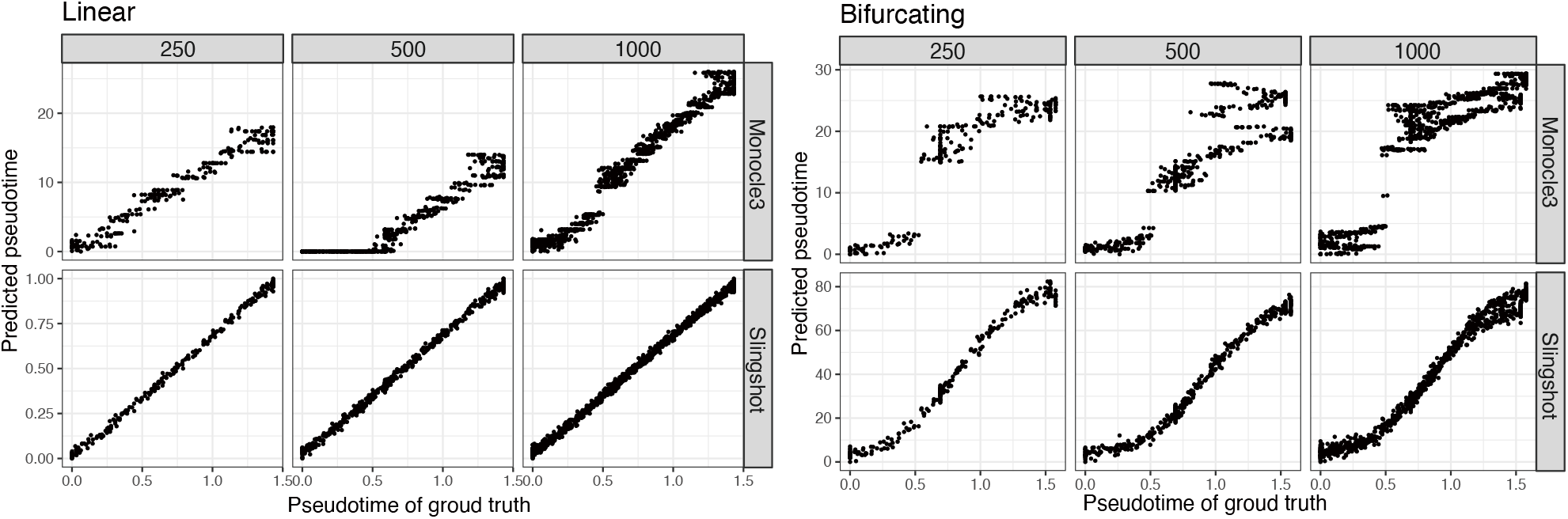
Correlation between the correct and predicted pseudotime in the simulation data. The numbers above the boxes indicate the number of cells. The trajectory inference is shown to the right of the box. Slingshot generally calculates accurate values, but Monocle3 produces a gap in some conditions.

**Fig. 7.**
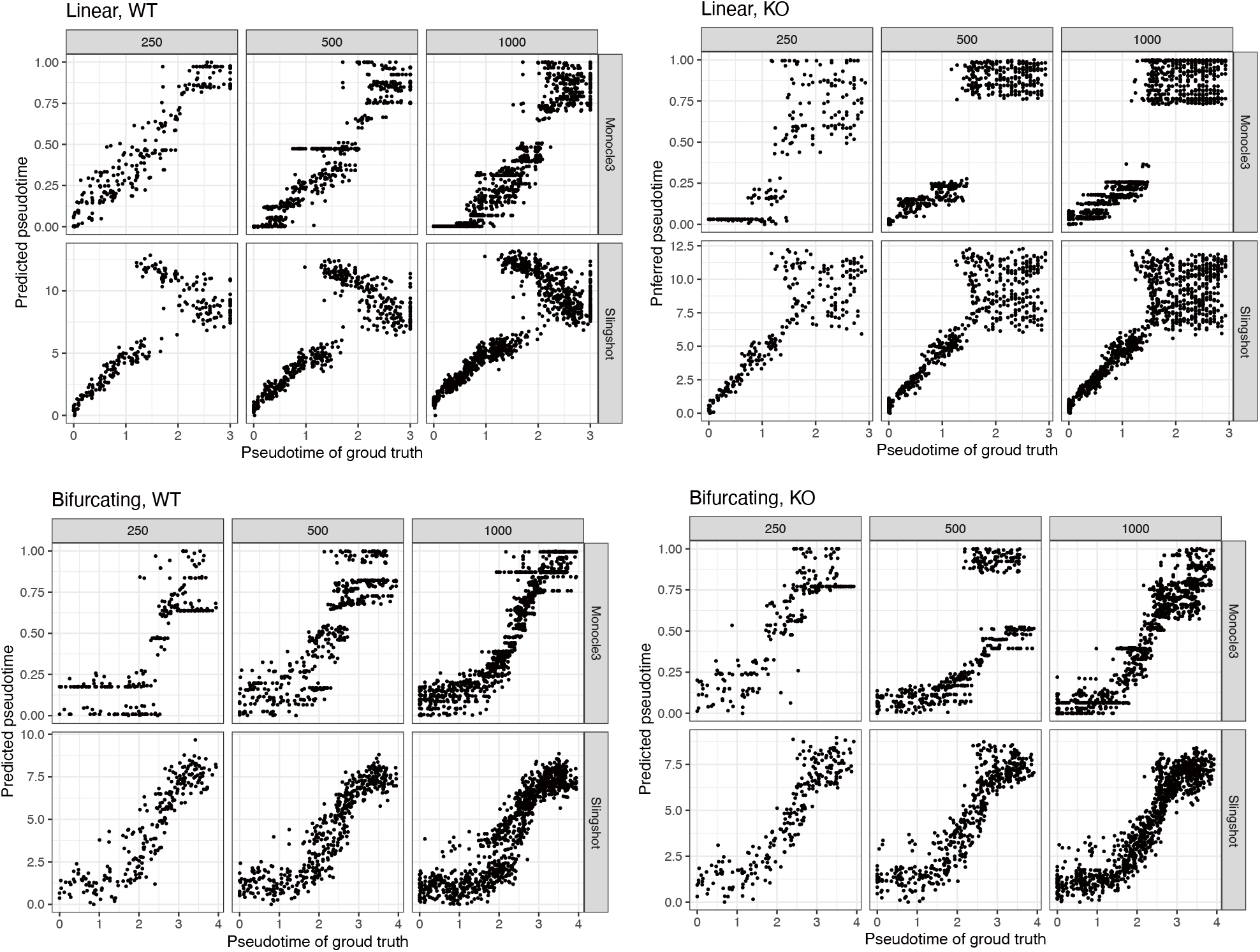
Correlation between the correct and predicted pseudotime in the simulation data. The numbers above the boxes indicate the number of cells. The trajectory inference is shown to the right of the box. Since there are two trajectories, WT and KO, the difference from ground truth is shown for each. Slingshot generally calculates accurate values, but Monocle3 produces a gap in some conditions.

A potential limitation of this study is that the benchmark was performed using a scRNA-seq data simulator rather than real experimental data. Although simulated data were used because the data from the experiments did not have the correct labels, they may have too little noise compared to the real data or may not mimic the real data. However, the issue in this study is not only the accuracy but also the introduction of frequency-domain analysis into trajectory analysis and its compatibility in any trajectory. Future comparisons of DEG analysis algorithms under more realistic conditions will be conducted using simulators and comparison strategies.

## Conclusion

In this study, we introduced a Lomb-Scargle periodogram-based algorithm to identify DEGs associated with pseudotime in trajectory analysis. The algorithm is capable of analyzing any inferred trajectory, including tree structures with multiple branching points, leading to diverse cell types. We validated this approach using simulated data and real datasets, and our results showed that our approach was superior when performing DEG analysis on complex structured trajectories. Our approach will contribute to gene characterization in trajectory analysis and help gain deeper biological insights.

## Author contributions statement

H.I.: Conceptualization, Data curation, Formal Analysis, Funding acquisition, Investigation, Methodology, Software, and Writing–original draft. M.H.: Project administration, Resources, Supervision, and Writing–review and editing.

## Acknowledgments

We are grateful to Ross Laidlaw, University of Glasgow, for providing the Trypanosoma dataset.

## Supplementary data

